# Distinct specificity of two pheromone G-protein coupled receptors, Map3 and Mam2, in fission yeast species

**DOI:** 10.1101/2020.07.22.215566

**Authors:** Taisuke Seike, Natsue Sakata, Chikashi Shimoda, Hironori Niki, Chikara Furusawa

## Abstract

Most sexually reproducing organisms have the ability to recognize individuals of the same species. In ascomycetes including yeasts, potential mating between cells of opposite mating-type depends on the molecular recognition of two peptidyl mating pheromones by their corresponding G-protein coupled receptors (GPCRs). Although such pheromone/receptor systems are likely to function in both mate choice and prezygotic isolation, very few studies have focus on the differences in pheromone/receptor system between mating types. The fission yeast *Schizosaccharomyces pombe* has two mating types (sexes), Plus (P) and Minus (M). Here we investigated the specificity of the two GPCRs, Mam2 and Map3, for their respective pheromones, P-factor and M-factor, in fission yeast. First, we switched GPCRs between *S. pombe* and the closely related species *Schizosaccharomyces octosporus*, which showed that SoMam2 (Mam2 of *S. octosporus*) is partially functional in *S. pombe*, whereas SoMap3 (Map3 of *S. octosporus*) is not interchangeable. Next, we swapped individual domains of Mam2 and Map3 with the respective domains in SoMam2 and SoMap3, which revealed differences between the receptors both in the intracellular regions that regulate the downstream signaling of pheromones and in the molecular recognition for pheromone binding. In particular, we demonstrated that two amino acid residues of Map3, F214 and F215, are essential for the specificity of M-factor recognition. Thus, the differences in these two GPCRs are likely to reflect the significantly distinct specificities of their respective pheromone/receptor systems; that is, the specificity of Map3 is more stringent than that of Mam2. We speculate that this sexual asymmetry might allow ascomycete fungi to generate novel prezygotic barriers within a population, while maintaining strong mate choice. Our genetic analyses also contribute to our understanding of the receptors that comprise the Class D GPCRs belonging to the fungal pheromone receptor family.

**Author summary:** Courtship signals play a key role in the fertilization processes of living beings from animals to microorganisms and, in particular, sex pheromones are involved in differentiating among species. On the other hand, changes in a pheromone/receptor system might also alter species-specificity during the selection of a mating partner, which is likely to facilitate prezygotic isolation. Here we demonstrate a distinct difference in the specificity of GPCRs for pheromones between cells of opposite mating-type in the fission yeast *Schizosaccharomyces pombe*, by conducting a comprehensive genetic analysis and comparison of these proteins between closely related fission yeast species. Our finding of sexual asymmetry due to the relative strictness of receptors in *S. pombe* suggests that it might be one of the driving forces behind the formation of new species. Such differences in pheromone/receptor interactions between the sexes may occur in a variety of life-forms such as insects and amphibians; hence, the findings of this study might be extended to other organisms. In addition, this study of *S. pombe* pheromone receptors sheds light on the functions of the unique Class D GPCRs.

## Introduction

Communication between cells is an essential feature of a life. In unicellular organisms such as yeasts, it is extremely important to find a compatible mating partner because zygote formation by cell fusion is a once-in-a-lifetime event [1, 2]. The specific ability of two cells of opposite mating-type to recognize each other is primarily determined by molecular recognition between a mating pheromone and its cognate receptor [3–6]. Such interactions are essential to differentiate between reproductively compatible and incompatible potential partners, and the specificity of the pheromone receptor regulates mate choice [3, 7–11]. In fact, our previous study demonstrated that genetic changes in a pheromone and its receptor genes trigger prezygotic isolation in the fission yeast *Schizosaccharomyces pombe* [12]. Those experimental data support the idea that even a simple pheromone/receptor system has significant ability to restrict gene flow. Therefore, the specificity of pheromone recognition must be stringently maintained among individuals of the same species.

However, a recent study on the budding yeast *Saccharomyces cerevisiae* found that yeast cells do not necessarily reject mates engineered to produce pheromones from other species, even if those species had been reproductively isolated from *S. cerevisiae* a long time ago [7]. Receptors might evolve to maximize reproductive success regardless of mate choice, potentially resulting in a relaxation of their specificity [14]. Such evolution might also provide an opportunity to mate with a related species producing different pheromones to promote genome diversity by genetic recombination. To date, however, only a few studies have investigated the specificity of pheromone receptors in yeast.

Most ascomycete fungi including yeasts produce two peptidyl mating pheromones: one is a farnesylated (and mostly *o*-methylated) lipid-peptide, whereas the other is an unmodified simple peptide. Notably, the distinct chemical modification states of the two pheromones are highly conserved among ascomycete fungi [3, 5]. Pheromone receptors belong to the family of G-protein coupled receptors (GPCRs). These receptors all share a common topology of seven α-helical transmembrane (TM) domains, including three extracellular loops (ELs) and three intracellular loops (ILs), and can be classified into distinct groups at the sequence level [15]. Fungal pheromone receptors are Class D GPCRs, which forms a distinct family phylogenetically distant from the well-studied rhodopsin-type (Class A) family. A study based on the primary structure of pheromone receptors of *S. cerevisiae* revealed that two types of pheromone receptor also exist in the genomes of other fungi [16]: one has a structure similar to that of a receptor for a lipid-peptide pheromone, whereas the other is homologous to a receptor for a simple peptide pheromone. The corresponding GPCRs for the two pheromones within each yeast species show no significant sequence similarity, consistent with the fact that the amino acid sequences of the two pheromone peptides bear no resemblance to each other.

*S. pombe* has two mating-types (sexes), termed Plus or P-cells (*h^+^*) and Minus or M-cells (*h*^−^) [17–19]. Under nitrogen-limited conditions, two haploid cells of the opposite mating-type form a diploid zygote, which then undergoes meiosis followed by the formation of four haploid spores within a cell [17, 18, 20–22]. M-cells secrete M-factor, a farnesylated and *o*-methylated nonapeptide, which is recognized by its corresponding GPCR Map3, expressed on the surface of P-cells [23, 24]. M-factor is encoded by three redundant genes, *mfm1^+^, mfm2^+^*, and *mfm3^+^* [23, 25]. These genes produce precursor polypeptides of 41–44 amino acids, each of which generates a single copy of the same M-factor, YTPKVPYMC^Far^-OCH_3_. M-factor is actively secreted by the specific ATP-binding cassette (ABC) transporter Mam1 [26]. By contrast, P-cells secrete P-factor, an unmodified peptide of 23 amino acids, which activates its cognate GPCR Mam2 on the surface of M-cells [27, 28]. P-factor is encoded by one gene, *map2^+^*, and is secreted by the exocytic secretary pathway rather than by active transport [28]. These pheromone signals stimulate cells of the opposite mating-type to commence a mating response. In *S. pombe*, both the Map3 and Mam2 receptors are coupled to the monomer form of the G-protein α-subunit Gpa1 [29, 30]. Activation of Gpa1 through the Map3 or Mam2 receptor transmits signals via a MAPK cascade consisting of Byr2 (MAPKKK), Byr1 (MAPKK), and Spk1 (MAPK) [30–32], thereby inducing the transcription of genes essential for mating [33, 34]. In other words, the signaling pathway downstream of the activated pheromone GPCRs is the same in P- and M-cells.

The homology between Map3 and Mam2 of *S. pombe* is low, a feature that is also observed between Ste3p (the receptor for the lipid-peptide pheromone a-factor) and Ste2p (the receptor for the simple peptide pheromone α-factor) of *S. cerevisiae*. Notably, sequence comparison between the pheromone receptors of *S. pombe* and *S. cerevisiae* shows that Map3 and Mam2 have significant homology with Ste3p and Ste2p, respectively, even though these yeast species are thought to have diverged between 300 million and 1 billion years ago [35]. For example, Mam2 and Ste2p share about 70% homology across the 5–7th TM domains [36]. This observation suggests that there is an evolutionary relationship between the two GPCRs for pheromones in fungi, presuming that the two sex receptors must have diverged early during evolution. Furthermore, this concerted evolution of the two pheromone GPCRs seems to have occurred in most ascomycete fungi.

We recently explored differences and similarities between genes encoding the two pheromones and their receptors in 150 wild *S. pombe* strains obtained from worldwide regions [13]. Sequencing analysis clearly demonstrated that there is no variation in the amino acid sequences of M-factor and Map3 in *S. pombe* strains, whereas P-factor and Mam2 show substantial diversity. Furthermore, all four P-factors of *S. octosporus* (SoP-factors) were partially recognized by Mam2 in *S. pombe* M-cells, but the single SoM-factor was not recognized by Map3 in *S. pombe* P-cells [13]. This implies that *S. pombe* is prezygotically isolated from *S. octosporus*, owing to a lack of compatibility of M-factor between the species. Recently, Greig’s group reported that Ste2p recognizes genetically divergent pheromones more easily as compared with Ste3p [7]. Their experimental data suggest that the specificity of Ste2p might be relaxed relative to that of Ste3p. Taken together, these observations suggest that the two GPCRs for pheromones in ascomycete fungi might have distinct specificities for pheromone recognition, but no direct evidence has been provided so far.

Here, we have carried out a series of investigations to document the specificity of the Map3 and Mam2 receptors in *S. pombe*. First, we investigated the mating frequency of *S. pombe* strains in which these pheromone receptors were replaced by those of the closely related species *S. octosporus*, which showed that *S. pombe* cells can mate with cells producing SoMam2 but not with cells producing SoMap3. Next, we used domain swapping between Map3 and SoMap3 to show that receptor regions including TM6 and EL3 are likely to be critical for M-factor recognition. Subsequent large-scale screening of mutant Map3 receptors that recognized SoM-factor in *S. pombe* cells identified F214 and F215 within the TM6 region as essential residues for interaction with M-factor peptide. Lastly, we showed that *S. pombe* cells carrying a gene set encoding SoM-factor and Map3 receptors with mutations of these residues instead of the wild-type gene set, can mate. Our extensive study demonstrates the distinct specificity of the two pheromone *S. pombe* receptors, which interact with a common G-protein to activate the same pathway, in *S. pombe*. We speculate that this very different recognition system between the sexes might allow ascomycete fungi to both maintain strong mate choice and facilitate outcrossing to increase diversity. We also discuss genetic information based on our experimental data to increase our understanding of the unique class D GPCRs.

## Results

### Prezygotic isolation between *S. pombe* and *S. octosporus* is probably due to the incompatibility of M-factor and Map3

To investigate the specificity of pheromone recognition by Map3 and Mam2, we first used genetic recombinant tests to examine whether the pheromone and receptor genes of *S. pombe* can be replaced with those of *S. octosporus*. In this test, two different drug-resistance markers were integrated into the chromosomes of heterothallic strains such that recombinants harboring both resistance markers would emerge if the two strains carrying different drug-resistance markers successfully mated. In other words, if no double-resistant colonies were observed, the two strains must be reproductively isolated. This sensitive recombinant test can detect extremely rare genetic recombination events.

We constructed *S. pombe* strains in which their own pheromone and receptor genes were replaced with the corresponding genes of *S. octosporus* (see Materials and Methods). Equal numbers of cells of heterothallic strains carrying different drug-resistance markers were mixed and spotted onto MEA medium to induce mating; after 2 days of incubation, an aliquot of this culture was spread onto YEA containing the appropriate drugs. The crosses between wild-type P-cells and M-cells secreting SoM-factors produced almost no double-resistant colonies (Fig 1), consistent with our previous data showing that SoM-factors are not recognized by Map3 [13]. Similarly, P-cells expressing SoMap3 were completely rejected by both wild-type M-cells and M-cells secreting SoM-factors (recombinant frequency, <10^−6^). By contrast, the crosses between wild-type M-cells and P-cells secreting SoP-factors generated abundant double-resistant colonies, and M-cells expressing SoMam2 were markedly accepted by both wild-type P-cells and P-cells secreting SoP-factors (Fig 1). These data indicated that the SoP-factor and SoMam2 gene are partially replaceable in *S. pombe*, but the SoM-factor and SoMap3 genes are not. Thus, *S. pombe* is likely to be prezygotically isolated from *S. octosporus*, at least in terms of the incompatibility of specificity between M-factor and Map3, rather than that between P-factor and Mam2.

**Fig 1.**
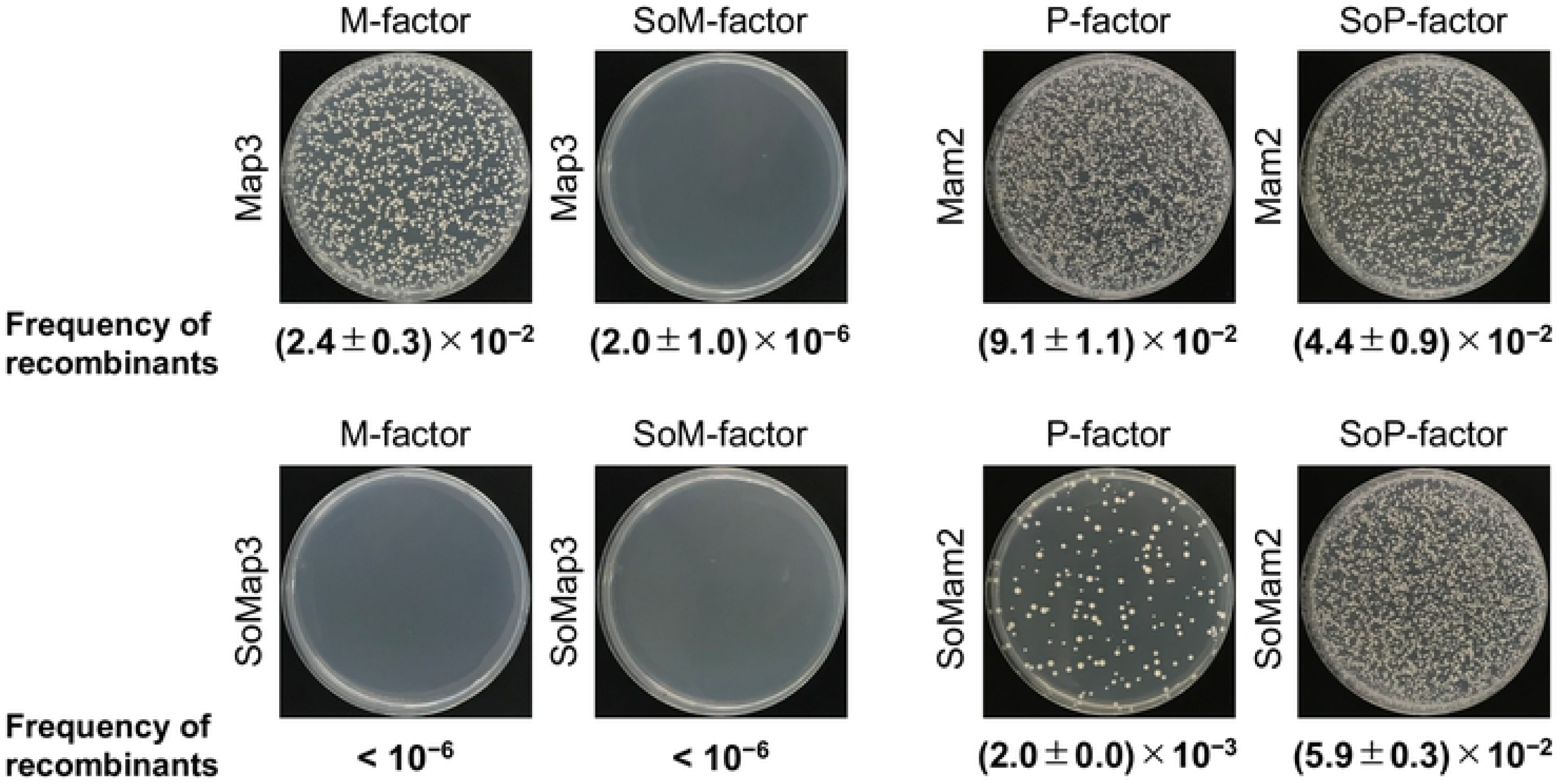
Prezygotic isolation test by recombinant frequency assay between two heterothallic *S. pombe* strains. Equal numbers of P- and M-cells were mixed and cultured on MEA medium for 2 days. The cells were then spread onto YEA medium containing 100 µg/ml of appropriate drugs. After 3 days of incubation, the plates were photographed and colony numbers were counted. The number of double-resistant colonies was normalized by the number of colonies on YEA plates without drugs. The assayed strains were TS620, M-strain secreting M-factor (kanMX6); TS9, M-strain secreting SoM-factor (kanMX6); TS26, P-strain expressing Map3 (hphMX6); and TS27, P-strain expressing SoMap3 (hphMX6) (Cross-I); and TS48, P-strain secreting P-factor (natMX6); TS49, P-strain secreting SoP-factor (natMX6); TS45, M-strain expressing Mam2 (hphMX6); and TS46, P-strain expressing SoMam2 (hphMX6) (Cross-II). Each assay was performed in triplicate; and the frequency of recombinants is reported as the mean ± SD.

### Differences in intracellular regions of the receptors partially prevent G-protein signaling

Fig 2A shows the structures of Map3 and Mam2 predicted by hydropathy analysis data [24, 27]. Although the amino acid sequences of Map3 and Mam2 both contain a high proportions of hydrophobic residues grouped into seven TM domains, there is no significant sequence similarity between them. The size of the proteins is similar, but Map3 and Mam2 differ markedly in the length of the N-terminal extracellular (EC) and C-terminal intracellular (IC) regions (Fig 2B and 2C). Overall, 240 residues are identical between Map3 (365 aa) and SoMap3 (363 aa; identity, 66%; Fig 2B), and 234 residues between Mam2 (348 aa) and SoMam2 (348 aa; identity, 67%; Fig 2C). Thus, homology comparison indicates that approximately two-thirds of the residues within these proteins are conserved between *S. pombe* and *S. octosporus*.

**Fig 2.**
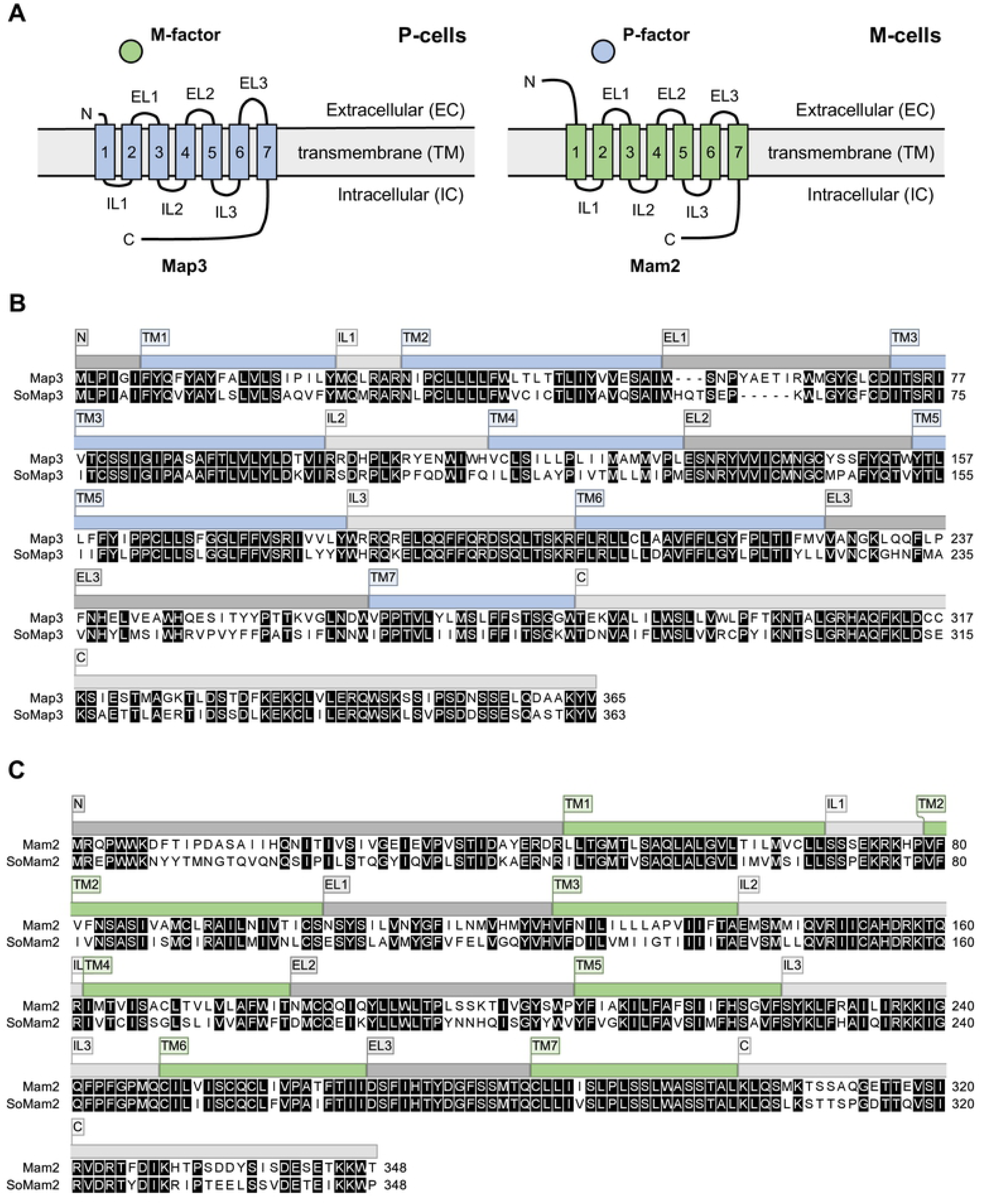
Comparison of the two pheromone GPCRs Map3 and Mam2. (A) Structures of Map3 (left) and Mam2 (right) predicted by hydropathy analysis. EL, extracellular loop; IL, intracellular loop; N, N-terminal tail, C, C-terminal tail. (B) Sequence alignment of the M-factor receptors Map3 (365 aa) and SoMap3 (363 aa). (C) Sequence alignment of the P-factor receptors Mam2 (348 aa) and SoMam2 (348 aa). Numerals to the right of each sequence indicate the length of the protein product. Identical amino acids are highlighted by a white letter on a black background. Protein domains are indicated above the sequence. The alignment was generated with CLC Genomics Workbench.

Expression of SoMap3 did not restore the mating ability of the *S. pombe map3Δ* strain, even in the presence of SoM-factor (Fig 1); therefore, we considered the possibility that SoMap3 might not efficiently activate *S. pombe* Gpa1, which transmits downstream pheromone signaling. To test this possibility, we first generated four Map3 chimeras, in which one IC domain was replaced with the corresponding sequence of *S. octosporus*, by domain-swapping using PCR with the appropriate primers (see Materials and Methods). The Map3 chimeras were then introduced into the *S. pombe map3Δ* strain and tested for mating frequency. Strains expressing each of the chimeric Map3 receptors did not show a significant decrease in mating frequency as compared with wild-type Map3 (Fig 3A). Next, therefore, we carried out stepwise replacement of the IC domains of Map3, which showed that mating frequency was greatly influenced by increasing changes of Map3; notably, the strain expressing the mutant Map3(IL1_2_3_C), in which all three ILs and the C-terminal region were replaced with the corresponding sequence of *S. octosporus*, produced less than 10% of the mating-cells of the strain expressing wild-type *map3^+^* (Fig 3B). These data suggested that intracellular regions of SoMap3 might not be compatible with Gpa1.

**Fig 3.**
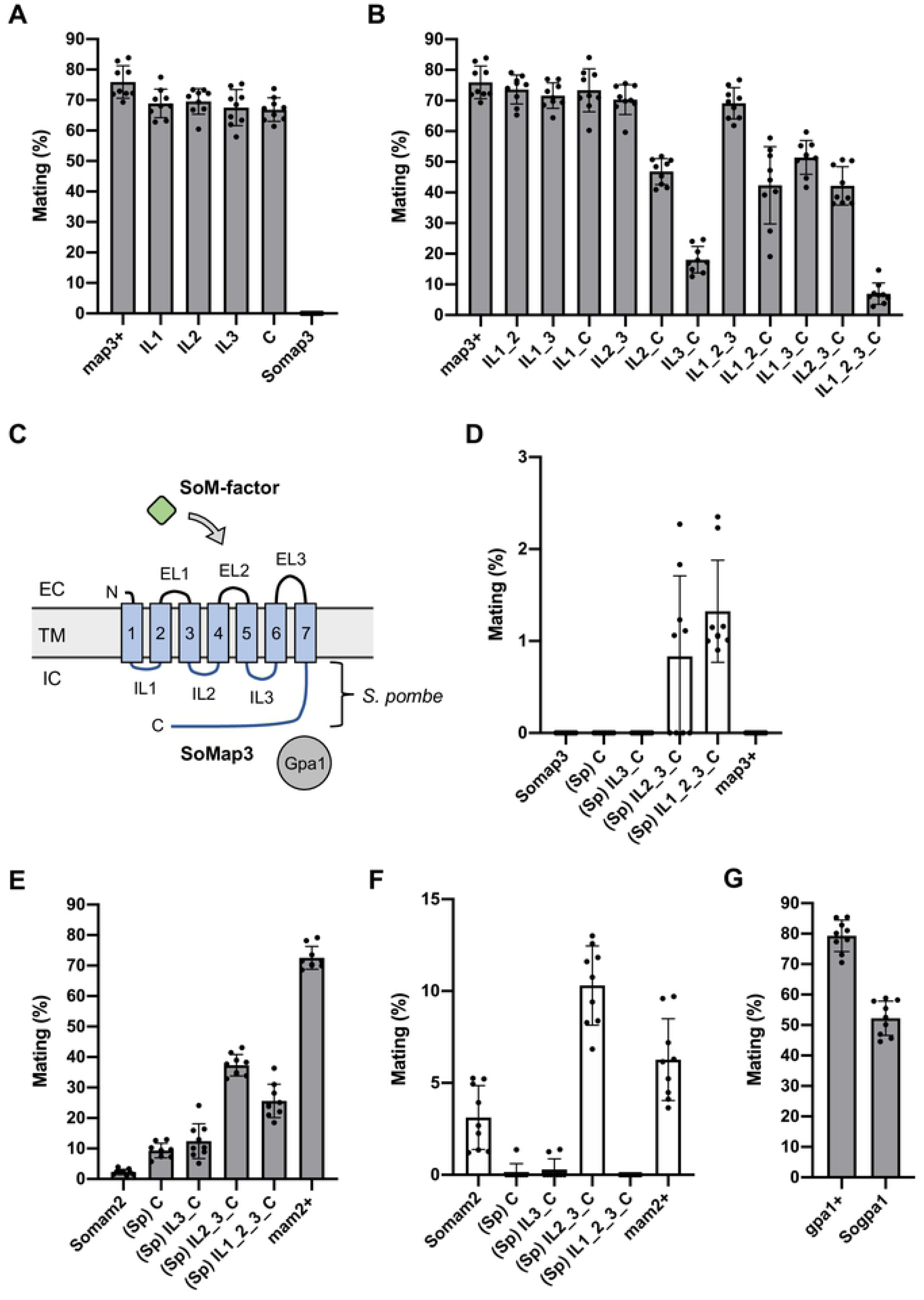
Genetic analysis of the intracellular regions of Map3 and Mam2 that are probably associated with Gpa1. (A) Mating frequency of homothallic M-factor-secreting strains each expressing a chimeric Map3 receptor in which one intracellular (IC) domain was replaced with the corresponding sequence of *S. octosporus*. (B) Mating frequency of homothallic M-factor-secreting strains each expressing a chimeric Map3 receptor constructed by the stepwise replacement of IC domains. (C) Predicted structure of SoMap3. EC, extracellular; TM, transmembrane; EL, extracellular loop; IL, intracellular loop; N, N-terminal tail, C, C-terminal tail. (D) Mating frequency of homothallic SoM-factor-secreting strains each expressing a chimeric SoMap3 in which the indicated IC domains were replaced with the corresponding sequence of *S. pombe*. (E) Mating frequency of homothallic P-factor-secreting strains each expressing a chimeric Mam2 in which one IC domain was replaced with the corresponding sequence of *S. octosporus*. (F) Mating frequency of homothallic SoP-factor-secreting strains each expressing a chimeric SoMam2 in which the indicated IC domains were replaced with the corresponding sequence of *S. pombe*. (G) Mating frequency of homothallic *Sogpa1* strain. At least 1,000 cells were examined for each strain. Data are the mean ± SD of nine independent samples. The numerical data are included in S6 Table.

Next, we considered whether a mutant SoMap3 receptor whose intracellular regions were replaced with those of Map3 would be able to activate Gpa1 on interaction with SoM-factors. To examine this hypothesis, chimeric SoMap3 receptors were introduced into an *S. pombe* strain secreting SoM-factors in the presence of Gpa1 (Fig 3C). The strain expressing the mutant SoMap3(IL1_2_3_C), in which all of three ILs and the C-terminal region were replaced with the corresponding sequence of *S. pombe*, were able to mate as expected (Fig 3D), although the mating frequency was extremely low (1.3 ± 0.6%).

We also found that replacement of the IC domains of SoMam2 with those of Mam2 improved compatibility between the receptor and Gpa1 to a greater extent as compared with Map3 (Fig 3E and 3F). In particular, the strain expressing the mutant SoMam2(IL2_3_C) showed improved mating capability with both a P-factor-secreting strain expressing the *map2^+^* gene (37.3 ± 3.5%) and an SoP-factor-secreting strain expressing the *map2^So^* gene (10.3 ± 2.2%) (Fig 3E and 3F). By contrast, the strain expressing the mutant SoMam2(IL1_2_3_C), in which all of three ILs and the C-terminal region were replaced with the corresponding sequence of *S. pombe*, showed lower mating frequencies as compared with SoMam2(IL2_3_C) (Fig 3E and 3F). These results suggest that the organization of the intracellular regions of the GPCRs might be essential for G-protein signaling. In addition, we found that a strain in which Gpa1 was replaced with SoGpa1 (Gpa1 of *S. octosporus*) showed a small decrease in mating frequency (Fig 3G). Taken together, these data imply that incompatibility between the intracellular regions of the two GPCRs (Map3 and Mam2) and Gpa1 is partially responsible for interrupting pheromone signaling.

### The sixth TM and third EC regions of Map3 are largely implicated in recognition of M-factor

We found that M-cells secreting SoM-factor barely mated with P-cells expressing SoMap3(IL1_2_3_C) (Fig 3D), which might be due to large structural changes of Map3 caused by replacing intracellular regions. Because the M-factor-binding site on Map3 has not been determined, we attempted to identify regions of Map3 involved in M-factor recognition. By replacing the EC and TM domains in turn with the corresponding sequences of *S. octosporus*, we generated 11 chimeric Map3 receptors. Most of the resultant strains expressing each of these chimeric receptors did not show a significant decrease in mating frequency; notably, however, the replacement of TM6 and EL3 led to a complete loss of mating ability (Fig 4A), meaning that these two domains might be essential for recognition of M-factor. We also swapped EC and TM domains of Mam2 with those of SoMam2; however, the replacement of these regions did not have a significant effect, except in the case of the N-terminal extracellular tail (Fig 4B). This observation may be related to differences in the molecular structures of Map3 and Mam2 that lead to the distinct specificity of each pheromone/receptor system.

**Fig 4.**
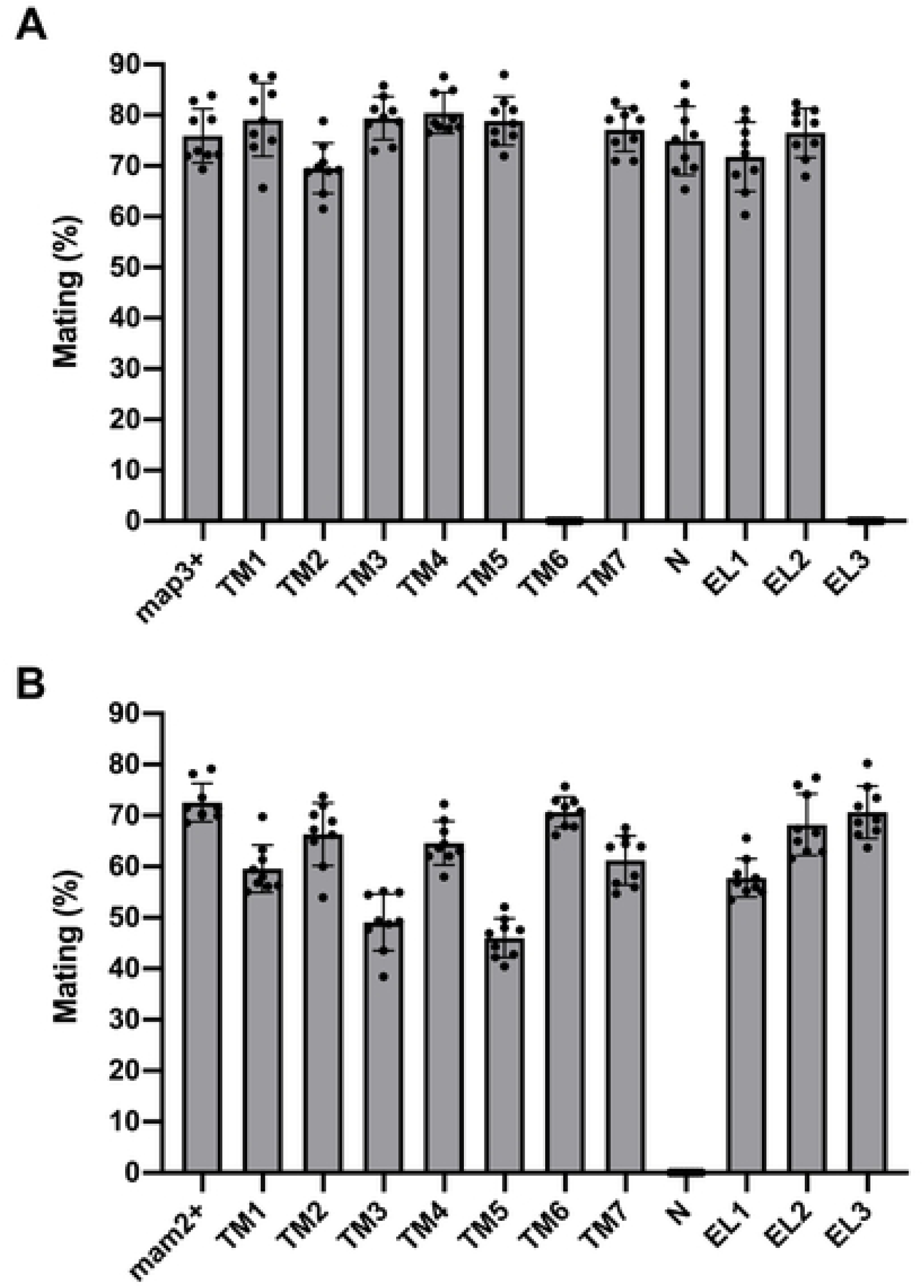
Genetic analysis of the extracellular regions and transmembrane domains of Map3 and Mam2. (A) Mating frequency of homothallic M-factor-secreting strains each expressing a chimeric Map3 in which one extracellular (EC) or transmembrane (TM) domains was replaced with the corresponding sequence of *S. octosporus*. (B) Mating frequency of homothallic P-factor-secreting strains each expressing a chimeric Mam2 in which one EC or TM domain was replaced with the corresponding sequence of *S. octosporus*. At least 1,000 cells were examined for each strain. Data are the mean ± SD of nine samples. The numerical data are included in S6 Table.

Next, we further subdivided the TM6 and EL3 region of Map3 (residues 204–264) into eight subdomains (TM6-1, TM6-2, and EL3-1–EL3-6), so that each subdomain contained 3–4 different amino acids between Map3 and SoMap3 (Fig 5A). Strains expressing the chimeric Map3 receptors were subjected to quantitation of mating frequency with M-factor secreting cells. Notably, replacement of the TM6-2 and EL3-1 domains led to a great reduction in recognition of M-factor by Map3 (Fig 5B). Based on these data, we presumed that residues critical for M-factor recognition are likely to lie within this region (residues 220–233). Therefore, we carried out site-directed mutagenesis of Map3 to create single substitutions of residues in this region (F224Y, M225L, V226L, A228V, G230C, L232G, and Q233H). Although one of these substitutions (Q233H) showed low mating frequency (4.1 ± 2.9%) (Fig 5C), we noticed that none of the Map3-Q233H, Map3(TM6-2_EL3-1), and Map3(TM6_EL3) mutants recognized SoM-factor (TS113, TS114, and TS117; sterile). Collectively, these data suggest that the region of Map3 corresponding to residues 204–264 must be important for M-factor recognition, but M-factor might bind to Map3 over a wider range. Nevertheless, this series of data indicate that the TM6 and EL3 domains of Map3 are largely implicated in M-factor recognition.

**Fig 5.**
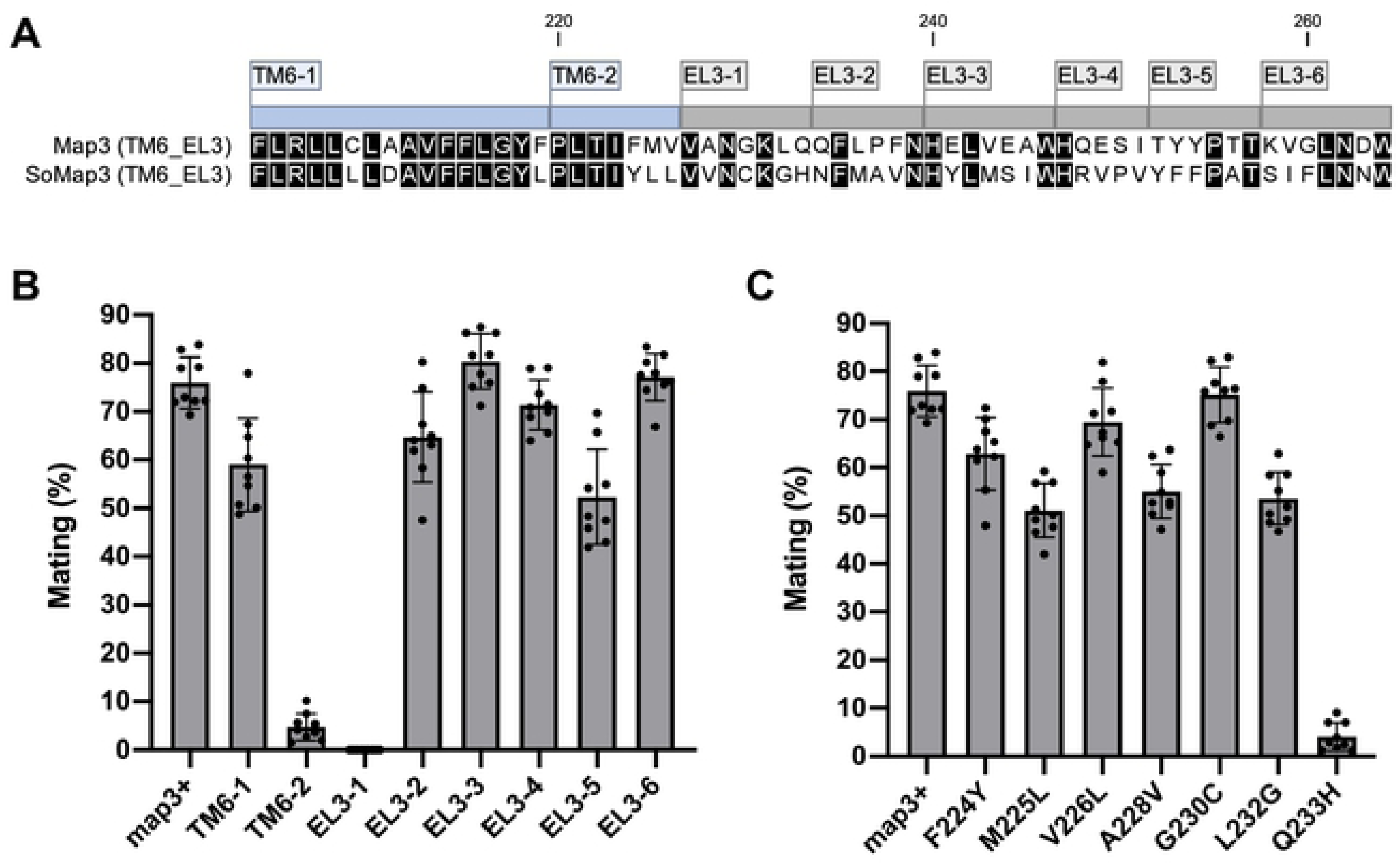
Genetic analysis of the region comprising residues 204–264 of Map3. (A) The 204–264-aa region of Map3 was subdivided into eight domains (TM6-1, TM6-2, and EL3-1–EL3-6), as indicated above the alignment sequence. (B) Mating frequency of homothallic M-factor-secreting strains each expressing a chimeric Map3 in which one of the eight domains was replaced. (C) Mating frequency of homothallic M-factor-secreting strains expressing Map3 with the indicated mutation. At least 1,000 cells were examined for each strain. Data are the mean ± SD of nine samples. The numerical data are included in S6 Table.

### Mutation of F214 and F215 in Map3 restores mating ability of *S. pombe* cells secreting SoM-factor

To find a mutant Map3 receptor that can efficiently recognize SoM-factor, we performed a random mutagenesis of the *map3* ORF (nucleotides 610–792, corresponding to residues 204–264) (Fig 6A). A high-quality *map3* mutant library constructed on a multicopy plasmid, pAHph-KS(map3^+^) (see Materials and Methods), was introduced into TS400, an *S. pombe* homothallic strain secreting SoM-factor. More than 10^5^ colonies on MEA were inspected for mating capability using efficient colony staining with iodine vapor, as described previously [12]. The large-scale screening yielded 232 putative mating-positive colonies, which were then microscopically inspected to eliminate false-positive clones (Fig 6B). Examination of mating capability identified 24 colonies in total. Plasmids were prepared from these colonies, and the nucleotide sequence of each *map3* gene was determined by Sanger sequencing, identifying 11 different mutant genes (Table 1). Next, each of these 11 suppressor *map3* mutant genes was integrated into the *map3^+^* locus on chromosome I, and strains showing significant mating ability, even in single-copy number, were selected. Ultimately, six genuine suppressor mutations were identified (Fig 6C).

**Fig 6.**
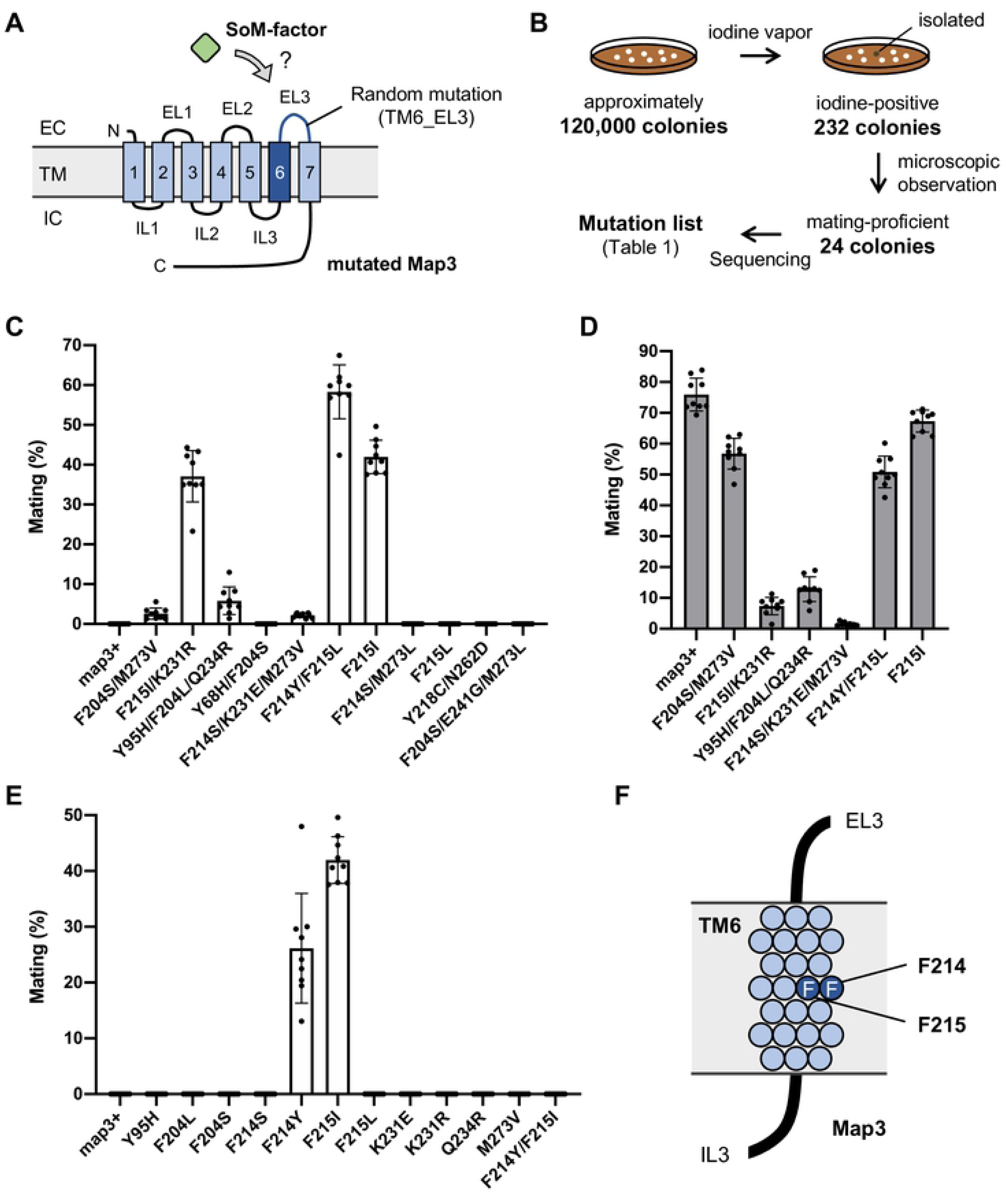
Identification of suppressor of Map3 that restore mating ability of homothallic SoM-factor-secreting strains. (A) Large-scale screening for mating-efficient colonies. Cells of a homothallic SoM-factor-secreting strain (TS400) carrying the *map3* mutant library introducing random mutations within the TM6/EL3 region were grown on MEA containing hygromycin B for 4 days at 30°C. (B) Approximately 1.2 × 10^5^ single colonies were inspected for sporulation capability by using efficient colony staining with iodine vapor. In total, 232 iodine-positive colonies were isolated and microscopically observed, resulting in 24 mating-proficient colonies. The *map3* genes contained in the plasmids prepared from these colonies were sequenced. All suppressor mutations identified are listed in Table 1. (C) Mating frequency of homothallic SoM-factor-secreting strains with one integrated copy of a suppressor *map3* gene. (D) Mating frequency of homothallic M-factor-secreting strains with one integrated copy of a suppressor *map3* gene that recognized SoM-factor. (E) Mating frequency of homothallic SoM-factor-secreting strains with one integrated copy of a *map3* gene with the indicated mutation. At least 1,000 cells were examined for each strain. Data are the mean ± SD of nine samples. The numerical data are included in S6 Table. (F) Positions of two potentially important residues (F214 and F215) for SoM-factor recognition by Map3 (blue circles). EL, extracellular loop; TM, transmembrane; IL, intracellular loop.

**Table 1.**
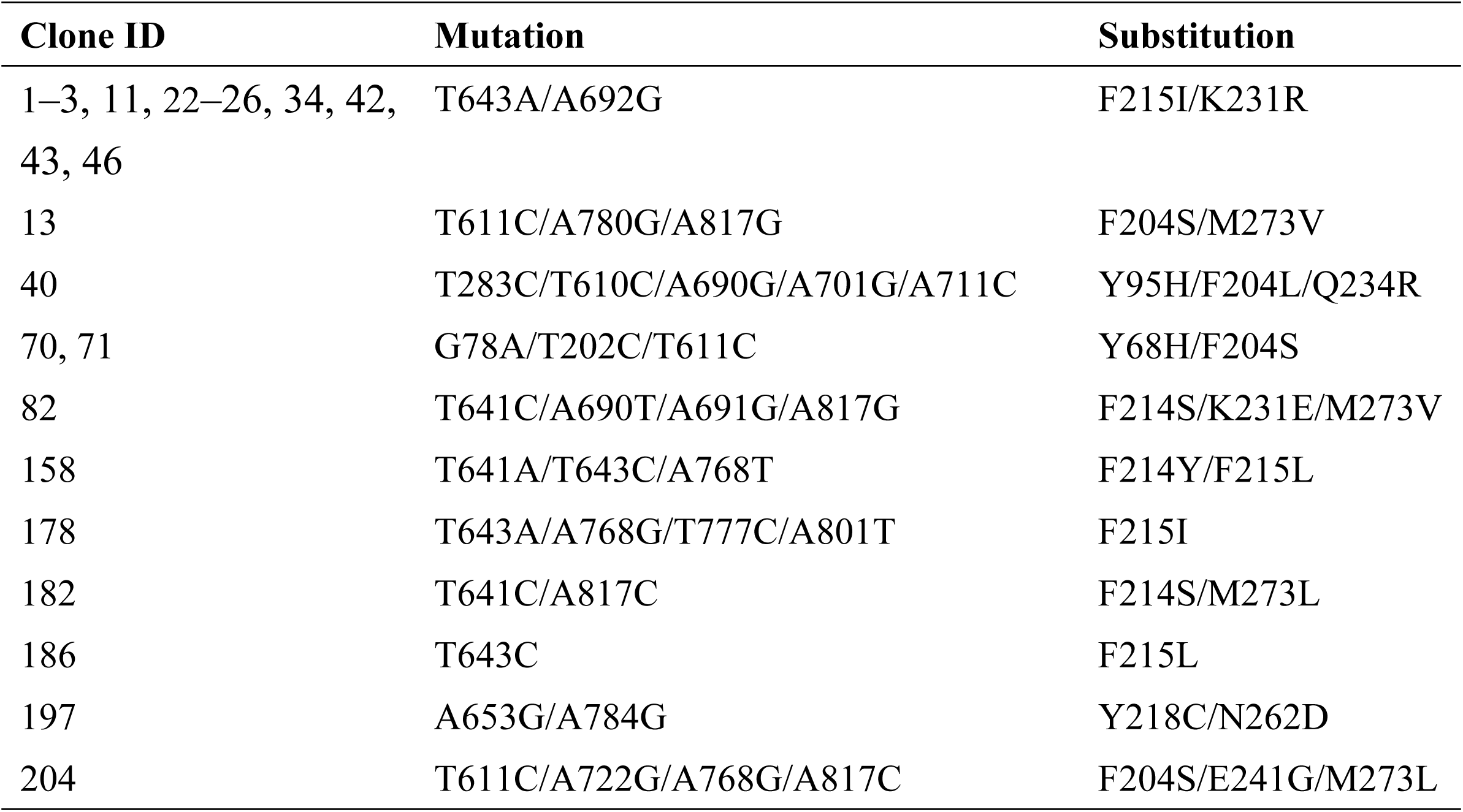
List of sterility-suppressing mutations of Map3 obtained by screening.

In particular, the strains expressing Map3-F215I/K231R, Map3-F214Y/F215L, or Map3-F215I showed fairly high mating frequency (37.1–58.3%; Fig 6C). We investigated whether these mutant Map3 receptors could also recognize M-factor by introducing them into the *map3Δ* strain secreting M-factor. We found that all six strains mated to some extent (Fig 6D), meaning that the mutant Map3 receptors were still able to recognize M-factor. Notably, the substitution K231R of Map3 seemed to prevent the efficient recognition of M-factor [Map3-F215I (67.3 ± 3.6%) vs Map3-F215I/K231R (7.4 ± 2.8%); Fig 6D].

Next, we focused on the individual amino acid substitutions found in the suppressor *map3* genes and determined whether one, two, or all three substitutions were required to provide beneficial effects for SoM-factor recognition. As shown in Fig 6E, F214Y and F215I were clearly the substitutions absolutely responsible for recognizing SoM-factor. However, a mutant Map3 harboring a double substitution of F214Y and F215I did not recognize SoM-factor (Fig 6E), suggesting that these two residues, F214 and F215, in the TM6 region are independently involved in the recognition of M-factor peptide (Fig 6F).

### Mutation of F214 and F215 triggers marked changes in the specificity of M-factor

The six identified suppressor substitutions of Map3 seemed to have low specificity for SoM-factor, because they retained the ability to mate with a strain secreting M-factor (Fig 6D). We therefore attempted to search for other mating-proficient Map3 receptors that could recognize SoM-factor. Two target residues (F214 and F215) were systematically substituted to generate 38 missense *map3* mutants (19 of the F214 mutants were previously reported by our group [12]), in which each of the two residues was substituted with the 19 other amino acids in turn. The 38 mutated *map3* genes were each integrated at the *map3^+^* locus of two receptor-less strains secreting either M-factor or SoM-factor. The mating frequency of the 76 (38 × 2) resultant mutant strains was then inspected by determining the frequency of mating-cells (Fig 7). Microscopic observations revealed that 10 of these substitutions (F214H, F214N, F214Y, F215A, F215G, F215I, F215T, F215V, F215W, and F215Y) showed some mating proficiency (>0%) via SoM-factor from M-cells. This comprehensive mating assay indicated that some intersex combinations of SoM-factor and mutant Map3 might be sufficiently functional even in *S. pombe*.

**Fig 7.**
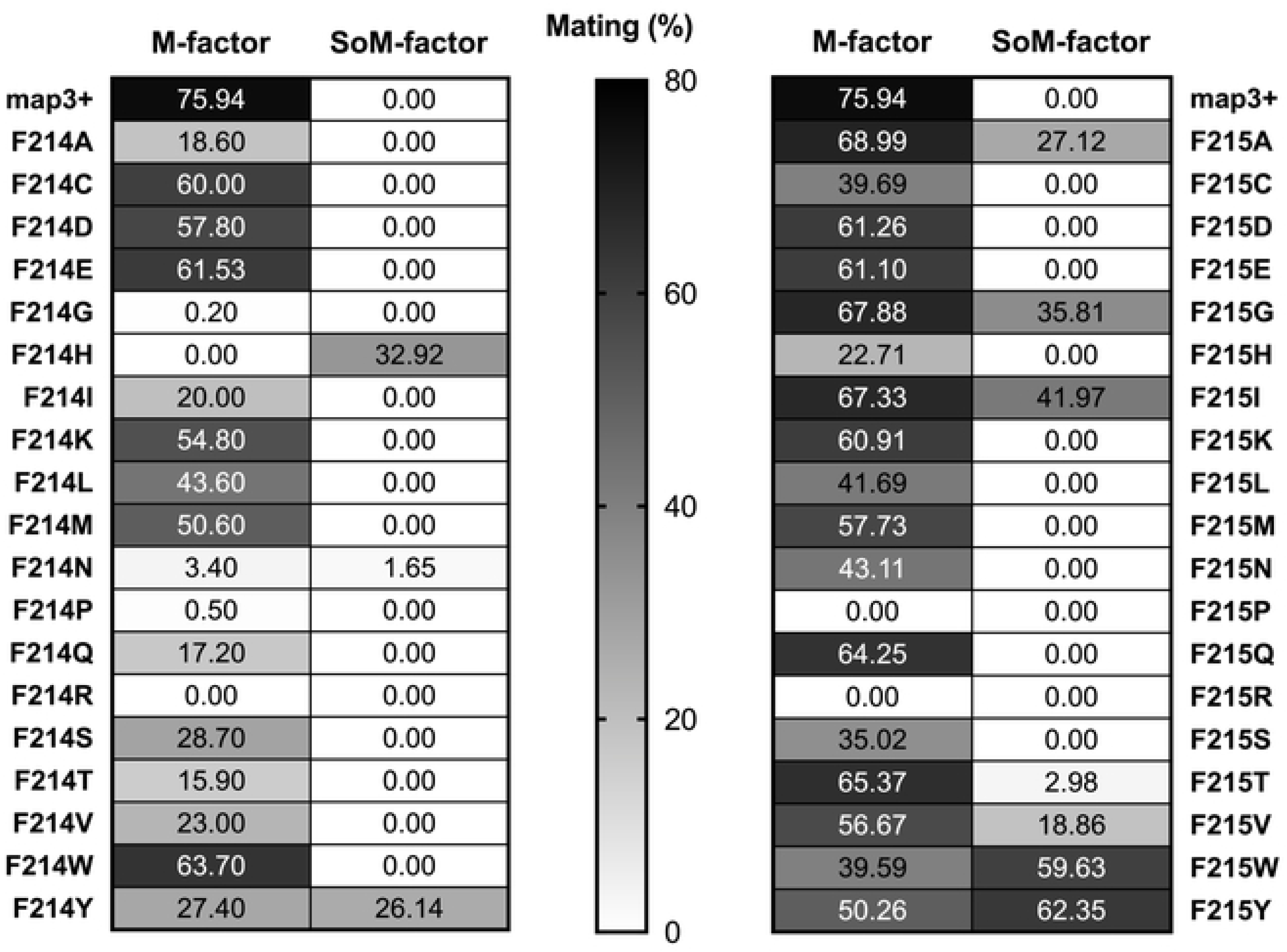
Mating ability of 76 *map3* missense mutants obtained after comprehensive site-directed mutagenesis of two key residues in Map3. Mating frequency of homothallic M-factor-secreting or SoM-factor-secreting strains, each expressing a missense Map3 receptor. Substitutions of the F214 and F215 residues of Map3 are indicated on the left. At least 1,000 cells were examined for each strain. Data are the mean ± SD of nine samples. The numerical data are included in S6 Table. Some of the values are extracted from our study [12]. Mating frequency is also indicated by the depth of shading, where darker shading indicates higher frequency.

To confirm these mating results and the improved specificity of Map3 with SoM-factor, the recognition levels of the wild-type and mutated Map3 receptors were compared *in vitro* by using synthetic M-factor peptide. Map4, a P-type-specific agglutinin, is expressed in P-cells dependent on M-factor signaling. Therefore, to assess whether the mutated Map3 receptors bind directly to SoM-factor peptides, we measured β-galactosidase activity under the *map4^+^* promoter (see Materials and methods). In brief, a map4^PRO^-lacZ fusion plasmid was introduced into heterothallic P-strains, which were grown in EMM2−N medium containing synthetic peptide (M-factor or SoM-factor) at 1 µM. After permeabilization of cells, β-galactosidase activity was assayed by using *o*-nitrophenyl-β-D-galactopyranoside (ONPG) as a substrate. Some of the tested Map3 receptors (>10% in Fig 7) recognized synthetic SoM-factor to a considerable extent (Table 2). As expected, two double mutants (F215I/K231R and F214Y/F215L), comprising substitutions identified from screening, gave a high level of Miller units in the presence of SoM-factor. We also found that the Map3-F215G showed the most efficient recognition, which was approximately 13-fold higher than that of wild-type Map3. In most cases, the mutant Map3 receptors maintained the ability to bind to M-factor, as well as SoM-factor peptides. This observation suggests that substitution of the F214 and F215 residues tends to relax the specificity of Map3, enabling it to recognize any M-factor. Taken together, these data demonstrate that a very few mutations of the TM6 region in Map3 bring about a substantial structural change facilitating the recognition of M-factors of other species, potentially allowing cross-reactions to occur between closely related species.

**Table 2.** Relative pheromone binding ability of mutated Map3 receptors.

| Substitution | Relative Miller units |  |
| --- | --- | --- |
|  | +M-factor (1μM) | +SoM-factor (1μM) |
| *map3*<sup>+</sup> | 100 | 100 |
| *Somap3* | 35.92 ± 12.68 | 48.80 ± 23.44 |
| F215I/K231R | 62.19 ± 5.51 | 432.11 ± 19.05 |
| F214Y/F215L | 198.37 ± 28.21 | 515.78 ± 66.25 |
| F214H | 4.29 ± 5.04 | 116.55 ± 50.43 |
| F214Y | 207.47 ± 9.20 | 130.17 ± 11.50 |
| F215A | 609.74 ± 95.28 | 480.73 ± 68.00 |
| F215G | 134.85 ± 17.38 | 1352.51 ± 257.90 |
| F215I | 120.58 ± 9.64 | 88.13 ± 19.80 |
| F215W | 265.32 ± 55.45 | 118.68 ± 37.22 |
| F215Y | 233.86 ± 34.55 | 128.76 ± 26.10 |

Miller units are expressed a relative value of the wild-type *map3*<sup>+</sup> gene (set at 100) after
subtraction of the control value (addition of methanol instead of M-factor peptide). $\beta$ - Galactosidase assays were performed in triplicate, and the mean ± SD is presented. The numerical data are included in S7 Table.

## Discussion

Genetic changes of a pheromone and its receptor can affect mate choice between mating partners, providing a prezygotic barrier within a population [7, 12]. In most ascomycetes fungi, each sex has one-to-one pair of a peptide pheromone and its corresponding receptor: as mentioned in the introduction, one is a lipid-peptide/GPCR; the other is a simple peptide/GPCR [3, 5]. There is no significant homology in amino acid sequence between the GPCRs, but their structures are highly conserved among fungal species. In this study, we investigated the specificity of Map3 and Mam2, the two pheromone receptors of *S. pombe*. Initially, we found that SoMap3 and SoM-factor are not functional in *S. pombe* cells, whereas SoMam2 and SoP-factors partially work (Fig 1). This asymmetry in pheromone recognition is really striking: it indicates that the two GPCRs vary in strictness of specificity for their pheromones, although they share the same downstream signaling pathway via the G-protein α-subunit Gpa1. A previous study in *S. cerevisiae* reported that the *STE3* and *STE2* genes, encoding Ste3p and Ste2p respectively, are differentially regulated at both the transcriptional and the post-transcriptional level by cryptic polyadenylation [37]. Thus, *S. cerevisiae* might adopt different strategies to regulate the gene expression of its two pheromone GPCRs. Even though the gene clusters that regulate the downstream signaling of the pheromone receptor may be similar in cells of opposite mating-type, they might vary from each other in terms of characteristics such as their expression and regulation, possibly resulting in the distinct pheromone specificity observed in yeasts.

Fungal pheromone receptors are class D GPCRs. The Mam2 receptor for P-factor contains a relatively longer N-terminal extracellular tail, reminiscent of Ste2p. Interestingly, both Mam2 and Ste2p form oligomers *in vivo* [29, 36]. In fact, some GPCRs have been shown to convert between the monomer and a dimeric complex, which might be a prerequisite for correct localization to the cell membrane and regulation [38, 39]. Oligomerization of Ste2p might be chiefly facilitated by interactions of its TM1 domain, while its N-terminus and other TM domains are important for complex formation [40–43]. The TM1 domain of Ste2p contains a (small)xxx(small) motif, where represents any amino acids, which is important for strong interactions among TM domains [44]. This motif comprises two small residues (typically glycine, but also alanine or serine) separated by three amino acids in the polypeptide chain, which physically places them on the same face of an α-helix [36]. Notably, Mam2 also contains six putative (small)xxx(small) motifs (G49-MTL-S53, S53-AGL-A57, A85-SIV-A89, S288-LPL-S292, S292-SLW-A296, and A296-SST-A300) in its TM1, TM2, and TM7 domains (S1 Fig). Lock *et al*. reported that mutations of these motifs in the TM1 domain affect oligomerization and reduce the function of Mam2 [36]. A comparison of Mam2 and SoMam2 shows that these motifs are highly conserved between *S. pombe* and *S. octosporus* (S1 Fig). Consistent with this, swapping the TM1, TM2, and TM7 domains between the two receptors had little effect on mating capability (Fig 4B); thus, it seems likely that these chimeric Map3 receptors formed a functional dimeric complex in *S. pombe*. By contrast, replacement of the N-terminal extracellular tail led to complete sterility (Fig 4B); therefore, the alteration of residues 1–45 might prevent the appropriate oligomerization of the receptor. In addition, SoMam2 seemed to interact poorly with intracellular proteins of *S. pombe*, but replacement of some ILs and the C-terminal region of SoMam2 with those of Mam2 led to a great improvement in mating capability (Fig 3E and 3F). Collectively, these experimental data suggest that the specificity of Mam2 for P-factors is not so stringent.

By contrast, the Map3 receptor contains a much shorter N-terminal extracellular tails, similar to Ste3p. There have been few studies of this type of GPCR, unlike Ste2p-like GPCRs [45, 46]. At present, therefore, oligomerization of Ste3p has not been confirmed, but there is only one (small)xxx(small) motif (A15-LVL-S19) in the TM1 domain of Map3 as far as we can tell (S2 Fig). In this study, systematic domain swapping of Map3 clearly showed that the TM6 and EL3 domains are critical for M-factor recognition (Fig 4A). Previously, we introduced random mutations into the whole region of the *map3^+^* gene to identify suppressor mutations that restored mating in sterile M-factor mutants [47], and successfully identified mutations mapped to three residues: F204, F214, and E249 [12]. Remarkably, all of these residues are located in the TM6 and EL3 domains. This observation supports the idea that the region comprising residues 204–264 is largely implicated in the specificity of M-factor. The F214 residue was identified as a suppressor mutation in our independent mutation studies (Table 1). Furthermore, our comprehensive mating assay showed that two consecutive phenylalanine residues, F214 and F215, affect binding to M-factor peptide (Fig 6F), with significant differences between the two residues. For example, F215 seemed to show a more gradually reduced specificity for M-factor as compared with F214 (Fig 7). In the *in vitro* pheromone binding assay, some of the mutant Map3 receptors in which F215 was substituted recognized SoM-factor at an unexpectedly high level (Table 2), suggesting that this substitution relaxes the specificity of Map3. We demonstrated that *S. pombe* P-cells expressing any of the single-mutant Map3 receptors (F214H, F214Y, F215A, F215G, F215I, F215W, and F215Y) were able to mate with *S. pombe* M-cells secreting SoM-factor (i.e., without wild-type M-factor or Map3) (Fig 6E and 7). These two residues, F214 and F215, are conserved among the genus *Schizosaccharomyces* (S2 Fig), but their mutation may disturb mate choice, potentially allowing *S. pombe* cells to form a diploid zygote by crossing with different species. Amino acid residues located in the TM6 domain are highly conserved, whereas the EL3 domain is largely varied among species (S2 Fig). Thus, the EL3 domain may be one of the M-factor-binding regions and the TM6 domain might facilitate structural changes at the pheromone-binding site. In this regard, we note that Q233 in the EL3, which affected mating efficiency (Fig 5C), is replaced by His (basic amino acid) in both *S. octosporus* and *Schizosaccharomyces cryophilus* and by Glu (acidic amino acid) in *Schizosaccharomyces japonicus* (S2 Fig). Lastly, replacement of IC domains in SoMap3 with those of Map3 barely altered the sterility of the *S. pombe* strain secreting SoM-factor (Fig 3D). This means that the IC regions are unlikely to be involved in M-factor recognition.

Further experiments are necessary to understand in detail the molecular recognition of pheromone peptides by GPCRs and the specificity of both the interaction between a pheromone and its receptor and between the receptor and downstream intracellular proteins. However, we have clearly demonstrated that the specificities of the Class D GPCRs Map3 and Mam2 differ significantly based on mating and recombinant assays using *S. pombe* strains expressing various chimeric receptor genes and pheromones. This difference is likely to be linked to the asymmetric diversification of M-factor and P-factor in nature [13]. Why would such an asymmetric setting be beneficial for yeast life? Although we have no explanations or working hypotheses for this finding at present, in future studies we plan to conduct simulation modelling of the evolutionary process to understand the effects of fixing pheromone communication in one direction. Fungal pheromone receptors might be the result of convergent evolution, shedding light on the pheromone signaling pathway and the functions of these uncharacterized GPCRs. Our study also showed that the TM6 and EL3 domains of Map3 are largely implicated in mate choice between *S. pombe* and *S. octosporus*. This finding might serve as a starting point for further mutagenesis studies to elucidate the structure–function relationship of the unique Class D GPCRs, as well as the molecular mechanism of prezygotic isolation caused by genetic changes of a pheromone/receptor system.

## Materials and methods

### Yeast strains, media, and culture conditions

The *S. pombe* and *S. octosporus* strains used in this study are listed in S1 Table. Standard methods at an incubation temperature of 30°C were used for growth, mating, and sporulation [17, 48, 49]. Yeast extract agar (YEA) medium supplemented with adenine sulfate (200 mg/ml), uracil (200 mg/ml), and L-leucine (100 mg/ml) was used for growth. Where appropriate, antibiotics (G418 disulfate [Nacalai Tesque, Kyoto, Japan], hygromycin B [Wako, Osaka, Japan], and nourseothricin [Cosmo Bio, Tokyo, Japan]; final concentration, 100 µg/ml) were added to YEA medium. Malt extract agar (MEA) medium was used for mating and sporulation. Synthetic dextrose medium was used to obtain auxotrophic mutants of *S. pombe*. Edinburgh minimal media (EMM2+N and EMM2−N) were used for growth in the β-galactosidase assay.

### Construction of plasmids carrying various M-factor-coding genes

The plasmids and oligo-primers used in this study are listed in S2 Table and S3 Table, respectively. All PCRs were performed by using KOD FX Neo DNA polymerase (TOYOBO, Tokyo, Japan).

To construct pTS271, the *mfm2^+^* gene (∼3.0 kb) containing the promoter and terminator regions was amplified from *S. pombe* (L968) genomic DNA using primer set oTS703/oTS704. Using Gibson Assembly (New England Biolabs, Ipswich, MA, USA), the DNA fragment was fused to a linearized vector derived from pFA6a-kanMX6 [50], which was prepared by inverse PCR using primer set oTS35/oTS36. Next, pTS274 was constructed by inverse PCR using pTS271 and primer set oTS707/oTS708. The amplified PCR product was treated with *Dpn*I, 5′-phosphorylated by T4 polynucleotide kinase (TaKaRa, Shiga, Japan), self-ligated by T4 ligase (TaKaRa, Shiga, Japan), and then introduced into *Escherichia coli* DH5α competent cells (New England Biolabs, Ipswich, MA, USA). The resulting plasmid was integrated into a recipient *S. pombe* strain after restriction with *Afl*II near the center of the promoter region.

To construct pTS272, the *mfm3^+^* gene (∼3.0 kb) containing promoter and terminator regions was amplified from L968 genomic DNA using primer set oTS705/oTS706. Using Gibson Assembly, the DNA fragment was fused to a linearized vector derived from pFA6a-natMX6 [50], which was prepared by inverse PCR using primer set oTS35/oTS36. Next, pTS275 was constructed by inverse PCR using pTS272 and primer set oTS709/oTS710. The resulting plasmid was integrated into a recipient *S. pombe* strain after restriction with *Afe*I near the center of the promoter region.

### Construction of plasmids carrying chimeric receptor genes

To construct chimeric receptor genes, we used a previously created plasmid, pFA6a-hphMX6(map3^+^) [12], which we renamed pTS185 (S2 Table). To construct pTS8, the *Somap3* (*map3^+^* of *S. octosporus*) ORF (1,092 bp) was amplified from *S. octosporus* (yFS286) genomic DNA using primer set oTS65/oTS66. Using Gibson Assembly, the DNA fragment was fused to a linearized vector derived from pTS185, which was prepared by inverse PCR using primer set oTS63/oTS64.

To construct plasmids carrying a chimeric *map3* gene, first, the corresponding region of *Somap3* was amplified by PCR using the appropriate primer set (S4 Table). Next, DNA replication using the amplified DNA fragment and pTS185 as a template was performed to generate each chimeric *map3* gene. After replication, the reaction mixture was treated with *Dpn*I to digest the template plasmid, which was transformed into DH5α. Each prepared chimeric region was confirmed by sequencing the recovered plasmid. Plasmids pTS19–pTS26, pTS42–pTS45, and pTS190–pTS199 were constructed by this method.

To replace the C-terminal region of *map3^+^* (nucleotides 850–1,098) with the corresponding region of *Somap3*, nucleotides 844–1,092 of *Somap3* were amplified from yFS286 genomic DNA using primer set oTS167/oTS168. Using Gibson Assembly, the DNA fragment was fused to a linearized vector derived from pTS185, which was prepared by inverse PCR using primer set oTS169/oTS170 to produce pTS46. Stepwise PCRs (a combination of the above) were also carried out to construct plasmids pTS91–pTS101 carrying chimeric *map3* genes in which multiple inner membrane regions were replaced.

To replace the C-terminal region of *Somap3* (nucleotides 844–1,092) with the corresponding region of *map3^+^*, nucleotides 850–1,098 of *map3^+^* were amplified from L968 genomic DNA using primer set oTS63/oTS750. Using Gibson Assembly, the DNA fragment was fused to a linearized vector derived from pTS8, which was prepared by inverse PCR with primer set oTS748/oTS749 to produce pTS292. Stepwise PCRs were also carried out to construct plasmids pTS294, pTS298, and pTS299 carrying chimeric *Somap3* genes in which multiple inner membrane regions were replaced by using the appropriate primer set (S4 Table). All resulting plasmids were integrated into the genome of a recipient *S. pombe* strain after restriction with *Nru*I in the terminator region.

To construct pTS32, the *Somam2* (*mam2^+^* of *S. octosporus*) ORF (1,047 bp) was amplified from yFS286 genomic DNA using primer set oTS128/oTS129. Using Gibson Assembly, the DNA fragment was fused to a linearized vector derived from pTS14, which was prepared by inverse PCR using primer set oTS149/oTS150. To construct plasmids carrying a chimeric *mam2* gene, inverse PCR was carried out using pTS14 and the appropriate primer set (S4 Table). The amplified PCR product was treated with *Dpn*I, 5′-phosphorylated by T4 polynucleotide kinase, self-ligated by T4 ligase, and transformed into DH5α. Each introduced mutation was confirmed by sequencing the recovered plasmid. Plasmids pTS305–pTS314 and pTS583 were constructed by this method.

To replace the C-terminal region of *Somam2* (nucleotides 904–1,047) with the corresponding region of *mam2^+^*, inverse PCR was carried out using pTS32 and primer set oTS825/oTS826 to produce pTS320. Stepwise PCRs were also carried out to construct plasmids pTS321–pTS323 carrying a chimeric *Somam2* gene in which multiple inner membrane regions were replaced by using the appropriate primer sets (S4 Table). All plasmids obtained were integrated into a recipient *S. pombe* strain after restriction with *Afe*I in the terminator region.

### Construction of plasmids carrying other pheromone signal-related genes

To construct pTS51, the *gpa1^+^* gene (∼2.2 kb) containing promoter and terminator regions was amplified from L968 genomic DNA using primer set oTS185/oTS186. Using Gibson Assembly, the DNA fragment was fused to a linearized vector derived from pFA6a-natMX6, which was prepared by inverse PCR using primer set oTS35/oTS36. To construct pTS300, the *Sogpa1* (*gpa1^+^* of *S. octosporus*) ORF (1,218 bp) was amplified from yFS286 genomic DNA using primer set oTS335/oTS336. Using Gibson Assembly, the DNA fragment was fused to a linearized vector derived from pTS51, which was prepared by inverse PCR using primer set oTS333/oTS334. The resulting plasmid was integrated into a recipient *S. pombe* strain after restriction with *Mfe*I near the center of the promoter region.

### Construction of a *map3* mutant library

To construct plasmid pAHph-KS (named pTS302), the hphMX6 marker (∼1.5 kb) containing TEF promoter and terminator regions was amplified from pFA6a-hphMX6 [50] using primer set oTS797/oTS798. Using Gibson Assembly, the DNA fragment was fused to a linearized vector derived from the multicopy plasmid pAL-KS [51], which was prepared by inverse PCR using primer set oTS795/oTS796. An ∼4 kb fragment containing the *map3^+^-*coding region and promoter sequence was amplified from L968 genomic DNA using primer set oTS811/oTS812. Using Gibson Assembly, the DNA fragment was fused to a linearized pTS302 vector, which was prepared by inverse PCR using primer set oTS807/oTS808. Using the resultant plasmid (named pTS317) as a template and primer set oTS950/oTS951, nucleotides 610–792 of the *map3* ORF were subjected to random mutagenesis by using the Diversify PCR Random Mutagenesis Kit (Clontech Laboratories, Mountain View, CA, USA) in accordance with the manufacturer’s instructions for error-prone PCR. To obtain a mutation rate of 5.8 mutations per 1 kb by error-prone PCR, the following reaction conditions were used: 34 ul of PCR-grade water, 5 µl of Titanium *Taq* Buffer (10×), 4 µl of MnSO_4_ (8 mM), 3 µl of dGTP (2 mM), 1 µl of Diversify dNTP Mix (50×), 1 µl of primer mix (10 µM each), 1 µl of template DNA (1 ng/µl), and 1 µl of Titanium *Taq* Polymerase. Using Gibson Assembly, the purified PCR products were fused to a linearized vector derived from pTS317, which was prepared by inverse PCR using primer set oTS952/oTS953 to generate a mutant library comprising ∼1.5 × 10^4^ independent *E. coli* colonies (named pTS370).

### Large-scale screening for *map3* mutants that suppress a mating-defective strain secreting SoM-factors

The pTS370 mutant library was transformed into strain TS400, which secretes SoM-factors. The transformants were plated on MEA medium containing hygromycin B (100 µg/ml) and colonies were inspected for diploid cells containing four spores, indicating mating ability. The colonies were subjected to iodine vapor, which turns sporulation-proficient colonies brown [12, 52]. From a screen of ∼1.2 × 10^5^ colonies, 232 iodine-positive colonies were obtained. Microscopic inspection identified 24 colonies with asci. Plasmids were prepared from these colonies and the *map3*-coding region was sequenced by using primer set oTS157/oTS158 (see Table 1 for the identified suppressor mutations).

To exclude the possibility that the observed suppression of mating sterility was due to an increase in copy number of relevant *map3* genes, each suppressor *map3* gene was cloned into the integration vector pFA6a. The DNA fragment containing the mutant *map3* gene was then amplified using primer set oTS124/oTS125 and fused via Gibson Assembly to a linearized vector derived from pTS185, which was prepared by inverse PCR using primer set oTS63/oTS64. These single-copy suppressor strains were examined for recovery of the sterility of M-factor mutants.

### Site-directed mutagenesis of the *map3^+^* gene

Plasmids carrying a single amino acid substitution of *map3* were constructed by inverse PCR using pTS185 and the appropriate primer set (S5 Table). The amplified PCR product was treated with *Dpn*I treatment, 5′-phosphorylated by T4 polynucleotide kinase, ligated by T4 ligase, and transformed into DH5α. Each introduced mutation was confirmed by sequencing the recovered plasmid. Where appropriate, stepwise inverse PCRs were performed, using the appropriate plasmid as a template, to construct plasmids carrying more than two amino acid substitutions of *map3*.

### Quantitative analysis of hybrid formation by recombinant frequency

To estimate prezygotic isolation between two *S. pombe* strains, recombinant frequency was determined by a quantitative assay of genetic recombination. Heterothallic strains each carrying a chromosomal drug-resistance marker (kanMX6, hphMX6, or natMX6) were cultured on YEA medium overnight, and equal numbers of P- and M-cells were mixed in sterilized water to an OD_600_ of 5.0 (1 × 10^8^ cells/ml). A 30-µl aliquot of the suspension was spotted onto MEA medium and incubated for 2 days at 30°C. The mixed cells were allowed to mate, and the resulting hybrid diploids were subjected to sporulation. The cell suspension was serially diluted and spread on YEA medium containing different combination of antibiotics. The number of colonies was counted after 3 days of incubation at 30°C. The ratio of recombinant frequency was calculated as described previously [12, 13, 53, 54]. Each analysis was carried out in triplicate, and the mean ± standard deviation (SD) was calculated.

### Quantitation of mating frequency

*S. pombe* cells grown on YEA medium overnight were resuspended in sterilized water to OD_600_ of 5.0. A 30-µl aliquot of the suspension was spotted onto MEA medium and incubated for 2 days at 30°C. The number of cells was counted under a differential interference contrast microscope (BX53, Olympus, Tokyo, Japan). Cell types were classified into four groups: vegetative (non-mating) cells (V), zygotes (Z), asci (A) and free spores (S). Mating frequency was calculated by the following equation [12, 13, 47, 53–55]:

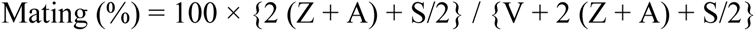

In all cases, cells were counted in nine randomly photographed digital images of each strain (>1,000 cells in total), and the mean ± SD was calculated (S6 Table).

### *In vitro* pheromone binding assay by a *map4* promoter–*lacZ* fusion gene

Given that P-type-specific agglutinin *map4^+^* mRNA expression is completely dependent on M-factor signaling [34] and the Map4 protein is expressed only in P-cells, expression of map4^PRO^–lacZ was assayed by β-galactosidase activity to assess whether the chimeric receptors recognized M-factor peptides. We used a previously created plasmid, pTA(map4-promoter:lacZ) [53], which we renamed pTS540 (S2 Table). First, pTS540 was introduced into a heterothallic P-type strain. For the assay, cells (OD_600_ 0.2) were grown in EMM2+N medium lacking leucine (to avoid loss of plasmid) for 24 hours, washed with EMM2−N medium three times, and then resuspended in EMM2−N medium at a cell density of 4 × 10^7^ cells/ml (OD_600_ 2.0). The cells were then treated with synthetic pheromone peptide (1 µM) [13] and incubated for exactly 24 hours with gentle shaking. The cell suspensions were centrifuged, and the cell pellets were resuspended in 1 ml Z-buffer (60mM Na_2_HPO_4_, 40 mM NaH_2_PO_4_, 10 mM KCl, 1mM MgSO_4_•7H_2_O, and 50 mM β-mercaptoethanol, pH7.0) and subjected to β-galactosidase assay using the sodium dodecyl sulfate (SDS)/chloroform cell permeabilization method, as previously described [56].

In brief, the diluted cells (OD_600_ 1.0) were permeabilized with 5 µl of chloroform:SDS (5:4 ratio) in 150 µl of Z-buffer for 5 min at 30°C, and then 30 µl of ONPG (4 mg/ml) was added as a substrate. The cell density of the cell suspension was measured, and then the A_420_ of the cell suspension was successively determined every 5 min by a microplate reader (Infinite 200 PRO, Tecan, Männedorf, Switzerland). The background activity measured in the absence of pheromone peptides (control) was subtracted from all readings, and the normalized β-galactosidase activity was expressed as Miller units [57]. Each β-galactosidase assay was performed in triplicate, and the mean ± SD was calculated (S7 Table).

### Data availability

All relevant data are included in the paper and its Supporting Information files.

## Acknowledgments

We thank the National Bio-Resource Project (NBRP), Japan, for providing yeast strains and plasmids. We are also sincerely grateful to Dr. Taro Nakamura of Osaka City University for helpful comments and all members of our laboratory for beneficial suggestions.

## Author contributions

Conceptualization: T Seike, C Shimoda, H Niki

Data curation: T Seike, N Sakata

Formal analysis: T Seike, N Sakata

Funding acquisition: T Seike

Supervision: C Furusawa

Writing – original draft: T Seike

Writing – review & editing: C Shimoda, H Niki, C Furusawa

## Supplementary Information

**S1 Fig. Sequence alignment of P-factor receptors among four species of the genus *Schizosaccharomyces***

Shown are Mam2 (348 aa), SoMam2 (Mam2 of *S. octosporus*, 348 aa), ScMam2 (Mam2 of *S. cryophilus*, 348 aa), and SjMam2 (Mam2 of *S. japonicus*, 348 aa). Numerals to the right of each sequence indicate the length of the protein product. Identical amino acids among the four species and highlighted by a white letter on a black background; amino acids conserved between at least two Mam2 receptors are highlighted by a white letter on a gray background. Protein domains are indicated above the sequence. Putative (small)xxx(small) motifs are indicated by a red frame. The incompatible region between Mam2 and SoMam2 identified from our domain swapping data is indicated by a blue frame. The alignment was generated with CLC Genomics Workbench.

**S2 Fig. Sequence alignment of M-factor receptors among four species of the genus *Schizosaccharomyces***

Shown are Map3 (365 aa), SoMap3 (Map3 of *S. octosporus*, 363 aa), ScMap3 (Map3 of *S. cryophilus*, 363 aa), and SjMap3 (Map3 of *S. japonicus*, 360 aa). Numerals to the right of each sequence indicate the length of the protein product. Identical amino acids among the four species highlighted by a white letter on a black background; amino acids conserved between at least two Map3 proteins are highlighted by a white letter on a gray background. Protein domains are indicated above the sequence. A putative (small)xxx(small) motif is indicated by the red frame. The incompatible region between Mam2 and SoMam2 identified from our domain swapping data is indicated by the blue frame. The alignment was generated with CLC Genomics Workbench.

**S1 Table. Strains used in this study**

**S2 Table. Plasmids used in this study**

**S3 Table. Primers used in this study**

**S4 Table. Primer sets for construction of plasmids carrying chimeric receptor genes**

**S5 Table. Mutation primer sets for site-directed mutagenesis of the *map3^+^* gene by inverse PCR**

**S6 Table. Numerical data for mating frequency experiments**

**S7 Table. Numerical data for β-galactosidase assay**

## References

1. Fisher RA. The genetical theory of natural selection. The genetical theory of natural selection. 2011. doi:10.5962/bhl.title.27468

2. Johansson BG, Jones TM. The role of chemical communication in mate choice. Biological Reviews. 2007. doi:10.1111/j.1469-185X.2007.00009.x

3. Gonalves-Sá J, Murray A. Asymmetry in sexual pheromones is not required for ascomycete mating. Curr Biol. 2011;21: 1337–1346. doi:10.1016/j.cub.2011.06.054

4. Bender A, Sprague GF. Pheromones and pheromone receptors are the primary determinants of mating specificity in the yeast Saccharomyces cerevisiae. Genetics. 1989.

5. Martin SH, Wingfield BD, Wingfield MJ, Steenkamp ET. Causes and consequences of variability in peptide mating pheromones of ascomycete fungi. Mol Biol Evol. 2011;28: 1987–2003. doi:10.1093/molbev/msr022

6. Jones SK, Bennett RJ. Fungal mating pheromones: Choreographing the dating game. Fungal Genetics and Biology. 2011. doi:10.1016/j.fgb.2011.04.001

7. Rogers DW, Denton JA, McConnell E, Greig D. Experimental Evolution of Species Recognition. Curr Biol. 2015;25: 1753–1758. doi:10.1016/j.cub.2015.05.023

8. McCullough J, Herskowitz I. Mating pheromones of Saccharomyces kluyveri: Pheromone interactions between Saccharomyces kluyveri and Saccharomyces cerevisiae. J Bacteriol. 1979. doi:10.1128/jb.138.1.146-154.1979

9. Burke D, Mendonca-Previato L, Ballou CE. Cell-cell recognition in yeast: Purification of Hansenula wingei 21-cell sexual agglutination factor and comparison of the factors from three genera. Proc Natl Acad Sci U S A. 1980. doi:10.1073/pnas.77.1.318

10. Hisatomi T, Yanagishima N, Sakurai A, Kobayashi H. Interspecific actions of α mating pheromones on the a mating-type cells of three Saccharomyces yeasts. Curr Genet. 1988. doi:10.1007/BF00365752

11. Leu JY, Murray AW. Experimental evolution of mating discrimination in budding yeast. Curr Biol. 2006. doi:10.1016/j.cub.2005.12.028

12. Seike T, Nakamura T, Shimoda C. Molecular coevolution of a sex pheromone and its receptor triggers reproductive isolation in Schizosaccharomyces pombe. Proc Natl Acad Sci. 2015;112: 4405–4410. doi:10.1073/pnas.1501661112

13. Seike T, Shimoda C, Niki H. Asymmetric diversification of mating pheromones in fission yeast. PLoS Biol. 2019;17: e3000101. doi:10.1371/journal.pbio.3000101

14. Cardé RT, Baker TC. Sexual Communication with Pheromones. Chemical Ecology of Insects. 1984. doi:10.1007/978-1-4899-3368-3_13

15. Vassilatis DK, Hohmann JG, Zeng H, Li F, Ranchalis JE, Mortrud MT, et al. The G protein-coupled receptor repertoires of human and mouse. Proc Natl Acad Sci U S A. 2003. doi:10.1073/pnas.0230374100

16. Shpakov AO. Serpentine type receptors and heterotrimeric G-proteins in yeasts: Structural-functional organization and molecular mechanisms of action. Journal of Evolutionary Biochemistry and Physiology. 2007. doi:10.1134/S0022093007010012

17. Gutz H, Heslot H, Leupold U LN. Handbook of genetics. King R, editor. Plenum Press, New York; 1974.

18. Egel R. Mating-Type Genes, Meiosis, and Sporulation. Molecular Biology of the Fission Yeast. 1989. doi:10.1016/b978-0-12-514085-0.50007-5

19. Egel R. Fission Yeast in General Genetics. The Molecular Biology of Schizosaccharomyces pombe. 2004. doi:10.1007/978-3-662-10360-9_1

20. Bresch C, Müller G, Egel R. Genes involved in meiosis and sporulation of a yeast. MGG Mol Gen Genet. 1968. doi:10.1007/BF00433721

21. Egel R. Physiological aspects of conjugation in fission yeast. Planta. 1971. doi:10.1007/BF00387025

22. Nielsen O. Mating-Type Control and Differentiation. The Molecular Biology of Schizosaccharomyces pombe. 2004. doi:10.1007/978-3-662-10360-9_18

23. Davey J. Mating pheromones of the fission yeast Schizosaccharomyces pombe: purification and structural characterization of M-factor and isolation and analysis of two genes encoding the pheromone. EMBO J. 1992. doi:10.1002/j.1460-2075.1992.tb05134.x

24. Tanaka K, Davey J, Imai Y, Yamamoto M. Schizosaccharomyces pombe map3+ encodes the putative M-factor receptor. Mol Cell Biol. 1993. doi:10.1128/mcb.13.1.80

25. Kjaerulff S, Davey J, Nielsen O. Analysis of the structural genes encoding M-factor in the fission yeast Schizosaccharomyces pombe: identification of a third gene, mfm3. Mol Cell Biol. 1994. doi:10.1128/mcb.14.6.3895

26. Christensen PU, Davey J, Nielsen O. The Schizosaccharomyces pombe mam1 gene encodes an ABC transporter mediating secretion of M-factor. Mol Gen Genet. 1997. doi:10.1007/s004380050493

27. Kitamura K, Shimoda C. The Schizosaccharomyces pombe mam2 gene encodes a putative pheromone receptor which has a significant homology with the Saccharomyces cerevisiae Ste2 protein. EMBO J. 1991. doi:10.1002/j.1460-2075.1991.tb04943.x

28. Imai Y, Yamamoto M. The fission yeast mating pheromone P-factor: Its molecular structure, gene structure, and ability to induce gene expression and G1 arrest in the mating partner. Genes Dev. 1994;8: 328–338. doi:10.1101/gad.8.3.328

29. Ladds G, Davis K, Das A, Davey J. A constitutively active GPCR retains its G protein specificity and the ability to form dimers. Mol Microbiol. 2005. doi:10.1111/j.1365-2958.2004.04394.x

30. Obara T, Nakafuku M, Yamamoto M, Kaziro Y. Isolation and characterization of a gene encoding a G-protein alpha subunit from Schizosaccharomyces pombe: involvement in mating and sporulation pathways. Proc Natl Acad Sci. 1991. doi:10.1073/pnas.88.13.5877

31. Xu HP, White M, Marcus S, Wigler M. Concerted action of RAS and G proteins in the sexual response pathways of Schizosaccharomyces pombe. Mol Cell Biol. 1994. doi:10.1128/mcb.14.1.50

32. Barr MM, Tu H, Van Aelst L, Wigler M. Identification of Ste4 as a potential regulator of Byr2 in the sexual response pathway of Schizosaccharomyces pombe. Mol Cell Biol. 1996. doi:10.1128/mcb.16.10.5597

33. Mata J, Bähler J. Global roles of Ste11p, cell type, and pheromone in the control of gene expression during early sexual differentiation in fission yeast. Proc Natl Acad Sci U S A. 2006. doi:10.1073/pnas.0603403103

34. Xue-Franzén Y, Kjærulff S, Holmberg C, Wright A, Nielsen O. Genomewide identification of pheromone-targeted transcription in fission yeast. BMC Genomics. 2006;7: 1–18. doi:10.1186/1471-2164-7-303

35. Painter RB, Nickoloff JA, Hoekstra MF. DNA Damage and Repair, Volume I: DNA Repair in Prokaryotes and Lower Eukaryotes. Radiat Res. 1998. doi:10.2307/3579915

36. Lock A, Forfar R, Weston C, Bowsher L, Uptond GJG, Reynolds CA, et al. One motif to bind them: A small-XXX-small motif affects transmembrane domain 1 oligomerization, function, localization, and cross-talk between two yeast GPCRs. Biochim Biophys Acta - Biomembr. 2014. doi:10.1016/j.bbamem.2014.08.019

37. Di Segni G, Gastaldi S, Zamboni M, Tocchini-Valentini GP. Yeast pheromone receptor genes STE2 and STE3 are differently regulated at the transcription and polyadenylation level. Proc Natl Acad Sci U S A. 2011. doi:10.1073/pnas.1114648108

38. Terrillon S, Bouvier M. Roles of G-protein-coupled receptor dimerization. EMBO Rep. 2004. doi:10.1038/sj.embor.7400052

39. Ward RJ, Xu TR, Milligan G. GPCR oligomerization and receptor trafficking. Methods in Enzymology. 2013. doi:10.1016/B978-0-12-391862-8.00004-1

40. Gehret AU, Bajaj A, Naider F, Dumont ME. Oligomerization of the yeast α-factor receptor: Implications for dominant negative effects of mutant receptors. J Biol Chem. 2006. doi:10.1074/jbc.M513642200

41. Kim H, Lee BK, Naider F, Becker JM. Identification of specific transmembrane residues and ligand-induced interface changes involved in homo-dimer formation of a yeast G protein-coupled receptor. Biochemistry. 2009. doi:10.1021/bi901291c

42. Overton MC, Blumer KJ. The extracellular N-terminal domain and transmembrane domains 1 and 2 mediate oligomerization of a yeast G protein-coupled receptor. J Biol Chem. 2002. doi:10.1074/jbc.M205368200

43. Wang HX, Konopka JB. Identification of amino acids at two dimer interface regions of the α-factor receptor (Ste2). Biochemistry. 2009. doi:10.1021/bi900424h

44. Russ WP, Engelman DM. The GxxxG motif: A framework for transmembrane helix-helix association. J Mol Biol. 2000. doi:10.1006/jmbi.1999.3489

45. Hagen DC, McCaffrey G, Sprague GF. Evidence the yeast STE3 gene encodes a receptor for the peptide pheromone a factor: gene sequence and implications for the structure of the presumed receptor. Proc Natl Acad Sci. 1986. doi:10.1073/pnas.83.5.1418

46. Boone C, Davis NG, Sprague GF. Mutations that alter the third cytoplasmic loop of the a-factor receptor lead to a constitutive and hypersensitive phenotype. Proc Natl Acad Sci U S A. 1993. doi:10.1073/pnas.90.21.9921

47. Seike T, Yamagishi Y, Iio H, Nakamura T, Shimoda C. Remarkably simple sequence requirement of the M-factor pheromone of Schizosaccharomyces pombe. Genetics. 2012;191: 815–825. doi:10.1534/genetics.112.140483

48. Egel R, Egel-Mitani M. Premeiotic DNA synthesis in fission yeast. Exp Cell Res. 1974. doi:10.1016/0014-4827(74)90626-0

49. Moreno S, Klar A, Nurse P. Molecular genetic analysis of fission yeast Schizosaccharomyces pombe. Methods Enzymol. 1991. doi:10.1016/0076-6879(91)94059-L

50. Hentges P, Van Driessche B, Tafforeau L, Vandenhaute J, Carr AM. Three novel antibiotic marker cassettes for gene disruption and marker switching in Schizosaccharomyces pombe. Yeast. 2005. doi:10.1002/yea.1291

51. Yazawa H, Ogiso M, Kumagai H, Uemura H. Suppression of ricinoleic acid toxicity by ptl2 overexpression in fission yeast Schizosaccharomyces pombe. Appl Microbiol Biotechnol. 2014. doi:10.1007/s00253-014-6006-y

52. R E. Fission yeast in general genetics. In: R E, editor. The Molecular Biology of Schizosaccharomyces pombe. Springer Berlin Heidelberg; 1974. pp. 1–12.

53. Seike T, Nakamura T, Shimoda C. Distal and Proximal Actions of Peptide Pheromone M-Factor Control Different Conjugation Steps in Fission Yeast. PLoS One. 2013;8: 1–15. doi:10.1371/journal.pone.0069491

54. Seike T, Maekawa H, Nakamura T, Shimoda C. The asymmetric chemical structures of two mating pheromones reflect their differential roles in mating of fission yeast. J Cell Sci. 2019. doi:10.1242/jcs.230722

55. Seike T, Niki H. Mating response and construction of heterothallic strains of the fission yeast Schizosaccharomyces octosporus. FEMS Yeast Res. 2017;17: 1–14. doi:10.1093/femsyr/fox045

56. Guarente L. Yeast Promoters and lacZ Fusions Designed to Study Expression of Cloned Genes in Yeast. Methods Enzymol. 1983. doi:10.1016/0076-6879(83)01013-7

57. Miller J. Experiments in Molecular Biology. Cold Spring Harbor, NY: Cold Spring Harbor Laboratory Press; 1972.

